# Educational Environment and White Matter Development in Early Adolescence

**DOI:** 10.1101/2023.10.10.561784

**Authors:** Ethan Roy, Amandine Van Rinsveld, Pierre Nedelec, Adam Richie-Halford, Andreas M. Rauschecker, Leo P. Sugrue, Ariel Rokem, Bruce D. McCandliss, Jason D. Yeatman

**Affiliations:** Graduate School of Education, Stanford University, Stanford, CA, USA; Department of Radiology and Biomedical Imaging, University of California San Francisco, San Francisco, CA, USA; Division of Developmental-Behavioral Pediatrics, Stanford University, Stanford, CA, USA; Department of Psychology and eScience Institute, University of Washington, Seattle, WA, USA

## Abstract

Coarse measures of socioeconomic status, such as parental income or parental education, have been linked to differences in white matter development. However, these measures do not provide insight into specific aspects of an individual’s environment and how they relate to brain development. On the other hand, educational intervention studies have shown that changes in an individual’s educational context can drive measurable changes in their white matter. These studies, however, rarely consider socioeconomic factors in their results. In the present study, we examined the unique effect of educational opportunity on white matter development, even when controlling other known socioeconomic factors. To explore this question, we leveraged the rich demographic and neuroimaging data available in the ABCD study, as well the unique data-crosswalk between ABCD and the Stanford Education Data Archive (SEDA). We find that educational opportunity is related to accelerated white matter development, even when accounting for other socioeconomic factors, and that this relationship is most pronounced in white matter tracts associated with academic skills. These results suggest that the school a child attends has a measurable impact on brain development for years to come.

## Introduction

Students who attend high-quality schools demonstrate higher academic achievement both in terms of reading and math scores^1–4^, as well as long-term outcomes such as college admissions and social mobility^5–7^. A common suggestion in the scientific literature^8,9^, and popular press^10,11^, is that the relationship between educational opportunity and academic outcomes reflects the influence that high-quality educational experiences might exert on brain development. Despite the well-established links between school quality and academic achievement, the specific relationship between educational opportunity and brain development remains unexplored.

Past studies, however, have demonstrated a relationship between brain development and various non-academic socioeconomic and environmental factors^12^. For example, diffusion MRI has revealed that higher family income predicts differences in white matter properties in adulthood^13^ and that parental income moderates the relationship between cognitive flexibility and tissue properties across a range of white matter tracts^14^. Furthermore, the influence of genetic heritability on white matter structure has been shown to be higher for individuals from high income backgrounds^15^. Together these findings suggest that the financial resources available to an individual during childhood influence and interact with brain development in complex ways that are not fully understood.

Additionally, lower levels of parental education have been linked to differences in white matter structure across multiple fiber tracts, including those purportedly underlying academic skills, such as the left arcuate fasciulus, left superior longitudinal fasciculus (SLF) and left inferior longitudinal fasciculus (ILF)^16–18^. These studies offer a variety of different (and sometimes conflicting) accounts of the relationship between white matter, parental education, and cognitive skills, with some results finding a brain-behavior relationship in individuals with lower levels of parental education^17^ and others suggesting that the relationship between parental education and cognitive behaviors is completely mediated by white matter properties^16,18^. What none of these studies address is why parental education affects white matter structure. In other words, are differences in parental education a proxy for a variety of environmental factors (broadly encompassed by the construct of SES) that influence white matter development? Or are there specific aspects of a child’s environment that are responsible for the link between measures of SES and brain structure?

Typical measures of socioeconomic status (SES) do not elucidate the specific aspects of an individual’s environment that drive differences in white matter development. The hormone cortisol has been shown to mediate the relationship between stressful life experiences and white matter properties^19^, suggesting that environmental stress, which has been linked to aspects of SES^20^, plays a role in white matter development. Furthermore, other specific environmental factors, such as home language^21^, screen use^22^, early childhood nutrition^23,24^, and adverse childhood events^25,26^ have also been linked to differences in white matter properties. Together, these findings suggest that typical measures of SES, such as parental income or education, may act as proxies for a confluence of other environmental factors that directly impact white matter development.

However, across these studies examining the link between SES and white matter development, an individual’s academic environment has gone unexplored. Although traditional measures of SES are highly correlated with school achievement^27^, it remains unclear the extent to which the specific school, and more generally the educational environment, that a child ends up in influences white matter development above and beyond the myriad of correlated factors that are wrapped up in indices of SES.

It bears mentioning that educational intervention studies have demonstrated that educational experiences can drive changes in brain structure and function over remarkably short timescales. These studies have shown that a short-term, intensive reading intervention led to changes across a range of white matter tracts, including left arcuate and left ILF, and that these changes correspond to changes in reading skill^28–30^. In the domain of mathematics, intensive learning experiences have been shown to “normalize’’ functional activity in students with mathematical learning difficulties^31^ and participation in specific math curricula drives changes in neurotransmitter concentration in the middle frontal gyrus and predicts longitudinal changes in mathematical reasoning^32^.

Although these findings serve as a proof-of-concept that educational experience can shape brain development in a manner that facilitates the development of academic skills, the samples used in these studies included a small number of participants in an intervention setting and were not representative of the population at large^33^. Moreover, the intensive and highly controlled interventions employed by these studies are far from representative of the typical differences among American schools. Furthermore, these studies did not include measures of SES in their analyses; It is possible or even likely that interventions have variable effects on brain development and learning depending on a child’s sociodemographic background^34,35^.

Although careful recruitment strategies can lead to sociodeomgraphcally diverse study populations, it is nearly impossible to capture the vast range of educational experiences of students across the United States in a typical brain imaging study. Recent efforts to collect and share large-scale neuroimaging datasets^36–40^ have now opened the door for researchers to explore the interplay between brain development, cognitive skills, and environmental and demographic factors. The ongoing ABCD study^38^ is particularly well positioned to examine the relationship between school quality and brain development in a large and representative sample. This study is following a cohort of approximately 10,000 children from across the United States longitudinally to understand brain development throughout adolescence. In addition to neuroimaging data, the ABCD study collects rich demographic and behavioral data on each participant, including traditional measures of SES, household and neighborhood cohesion, and educational opportunity, as measured by the Stanford Education Data Archive (SEDA; see Methods for overview)^27^. This rich set of neuroimaging and demographic data presents the first opportunity to understand the relationship between brain development and the diversity of educational environments experienced by students across the United States.

In the current study, we test the hypothesis that differences in white matter development are related to the quality of an individual’s educational environment, while controlling for the multitude of factors indexed by traditional measures of SES. We first leverage the individual white matter tract data generated through automated-fiber quantification (AFQ)^41,42^ to test the hypothesis that educational opportunity relates to white matter development in specific tracts underlying academic skill, such as reading and math. We find that FA in the bilateral arcuate fasciculus, left posterior arcuate, and the corpus callosum are related to educational opportunity, even when controlling for other measures of socioeconomic status. We then train a brain-age model to test the hypothesis that educational opportunity relates to accelerated white matter development. This model suggests that an individual’s educational opportunity may influence white matter development throughout the brain, though this relationship may be more pronounced in white matter tracts associated with academic skills.

## Results

### Exploring the relationship between educational opportunity and white matter development

Intervention studies have shown that an individual’s academic environment can drive measurable changes in brain structure and function^28,29,31^. However, it remains unclear the extent to which educational opportunities uniquely influence brain development above and beyond other socioeconomic factors^12,16,19^. Here we operationalize educational opportunity based on data from the Stanford Educational Data Archive (SEDA)^27^. SEDA leverages standardized test scores from schools across the United States to calculate two indices of an individual’s educational environment. “Early educational opportunities” provided by parents, neighborhoods, pre-K, and early elementary school education^43^ are operationalized by the average 3rd grade test scores for a district (SEDA intercept). Year-to-year learning opportunities afforded by an individual school is operationalized by the average change in test scores for a school between grade 3rd and 8th (SEDA slope). Both measures are in z-score units and relative to national norms. Thus, a school with a SEDA intercept and slope of zero performs at the national average in terms of third grade test scores and in terms of how quickly students grow from year to year. A school with a SEDA intercept of -1 and slope of zero performs 1 standard deviation below the national average, and students progress at the average rate meaning the discrepancy in achievement is maintained throughout schooling. Because the ABCD study begins in 4th or 5th grade, SEDA intercept is the most relevant measure of the educational opportunity that a participant has experienced up until the first ABCD measurement.

As expected, SEDA intercept is highly correlated with traditional measures of socioeconomic status (Figure 1A). The relationships between these various indices of SES raise the possibility that measures such as parental education and household income could act as proxies for other factors, like educational opportunity, that might more directly influence brain development. To this end, we leverage the rich demographic and neuroimaging data available in the ABCD dataset^38^ to separate the unique contributions made by a student’s academic environment towards their white matter development, compared to other known socioeconomic factors.

**Figure 1:**
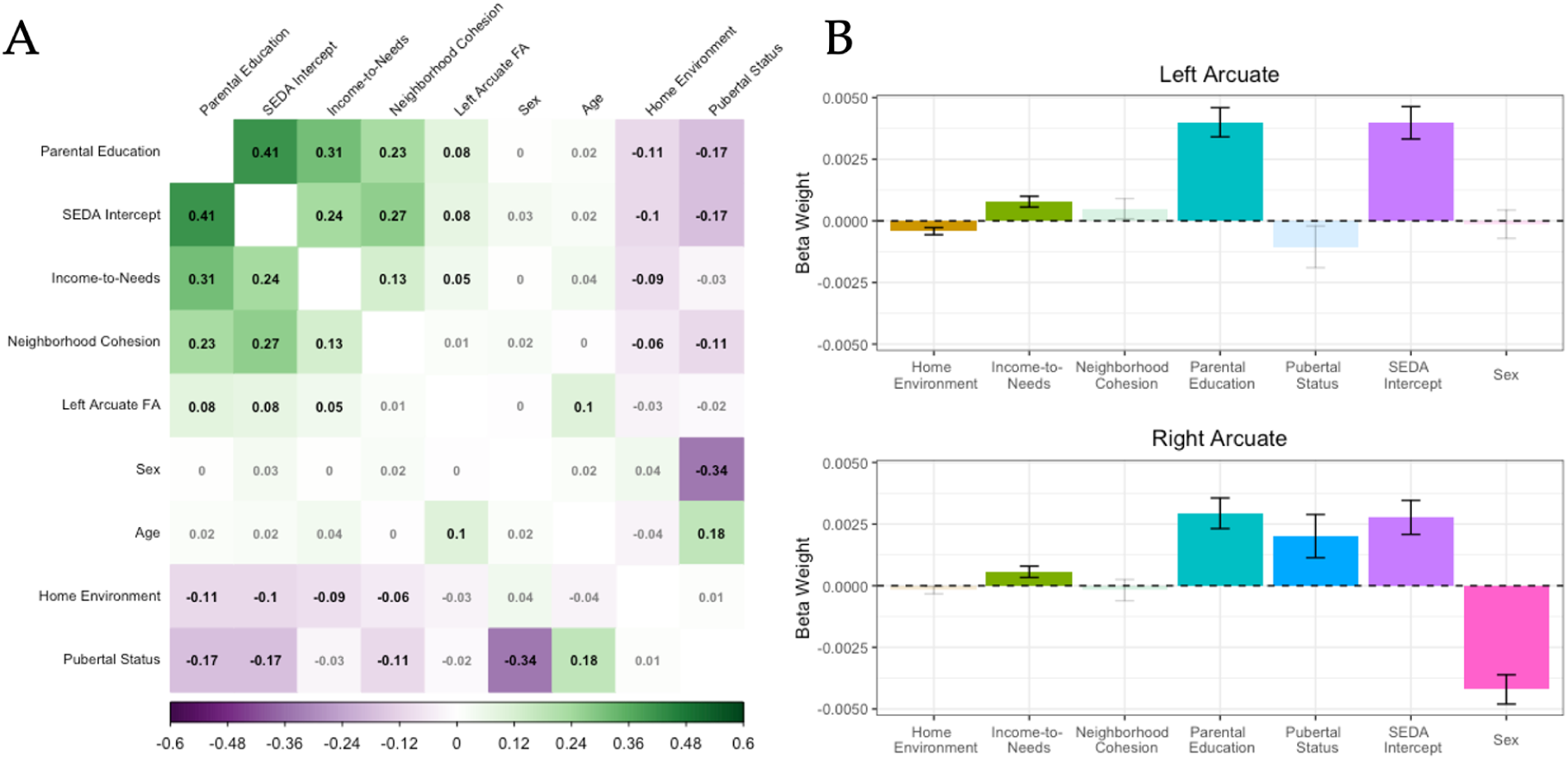
**A.** Correlation matrix illustrating the univariate relationships between mean FA in the left arcuate, SEDA intercept, and other demographic and socioeconomic factors. Coefficients in bold represent correlations where FDR-corrected p<0.05 .**B.** Beta-weights for linear mixed-effects models predicting mean FA in the left and right arcuate from a single predictor, specified on the x-axis. Each model included a random effects structure of family structure nested within scanner site. The colors of each bar denote each predictor variable. Error bars represent the standard error of each beta-coefficient. Bars that are bolded illustrate the beta-weights with FDR-corrected p<0.05.

#### Diffusion Properties of the Left Arcuate are Correlated with Socioeconomic Factors

Past studies have demonstrated a relationship between white matter development and socioeconomic factors, such as parental income, environmental stress and home context^14,19,21^. We began by attempting to replicate the results from past studies that observed univariate relationships between white matter and socioeconomic factors. We focus these initial analyses on the left arcuate, as past studies have shown this tract to relate to both reading skill and measures of SES^18,28,44^. We first calculated the correlation between mean FA in the left arcuate and a range of demographic and developmental factors, including age, pubertal status, sex, family cohesion, log-transformed income-to-needs ratio, parental education, SEDA intercept, and neighborhood cohesion (Figure 1A). This revealed small, yet significant correlations between mean FA in the left arcuate and parental education (*r* = 0.085; p_corrected_ < 0.001), income-to-needs ratio (*r* = 0.046 ; p_corrected_ = 0.019), SEDA intercept (*r* = 0.076; p_corrected_ < 0.001) and age (*r* = 0.096; p_corrected_ < 0.001).

However, these correlations do not account for individual and site-level random effects that might relate to white matter properties. To explore univariate relationships between FA in the left arcuate and demographic, developmental, and socioeconomic factors while controlling for these random effects, we constructed a series of linear mixed-effects models. These models included family structure nested within scanner site as random effects^45^ and either log-transformed income-to-needs ratio, parental education, family cohesion, neighborhood cohesion, sex, SEDA intercept or pubertal status as a sole fixed-effect predictor. As a control measure, we also fit the same sequence of models predicting FA in the right arcuate to assess whether these univariate relationships occur across the white matter or are specific to the left arcuate.

Similar to the correlation analyses, these models revealed significant relationships between mean FA in the left arcuate and parental education, SEDA intercept, and income-to-needs ratio (all p_corrected_ < 0.001; Figure 1B). Additionally, these models identified a slight yet significant relationship between home environment and mean FA in the left arcuate (B = -0.0004, p_corrected_ = 0.006) . Interestingly, the models predicting FA in the right arcuate also found significant relationships between mean FA and parental education, income-to-needs ratio, and SEDA intercept (all p_corrected_ < 0.05) as well as significant effects of pubertal status (B = 0.002, p_corrected_ = 0.023) and biological sex (B = -0.004, p_corrected_ < 0.001). Taken together, these models suggest that, in a univariate setting, FA and socioeconomic predictors, like income or parental education, relate to white matter properties.

#### The relationship between educational opportunity and tissue properties varies across the white matter

Although univariate analyses suggest a relationship between FA and socioeconomic factors, SES measures are also highly correlated with each other. Thus, the observed relationship between FA and any one index of SES may actually be driven by a separate, yet correlated measure. To test the hypothesis that there is a specific relationship between educational opportunity and white matter properties, while accounting for other developmental and socioeconomic effects, we modeled the relationship between SEDA intercept and tissue properties across all the white matter tracts identified by pyAFQ (see Methods of overview). Briefly, pyAFQ identifies 28 major white matter tracts and calculates diffusion properties at 100 nodes along the length of each tract (see Supplemental Figure 1 for overview of the tracts identified by pyAFQ). To account for scanner differences between the 21 ABCD study neuroimaging sites, we also performed ComBat harmonization^46–48^ across each node included in the pyAFQ outputs.

Based on previous research in smaller samples linking specific white matter tracts to academic skills, we hypothesized that educational opportunity would specifically relate to tissue properties in the the left arcuate fasciculus (ARC), left posterior arcuate fasciculus (pARC), bilateral inferior longitudinal fasciculus (ILF), bilateral superior longitudinal fasciculus (SLF), and uncinate fasciculus (UNC), as these tracts have all been previously implicated in academic skills, such as reading and arithmetic^49–52^. Within each tract, we then fit a linear mixed-effects model predicting mean harmonized fractional anisotropy (FA; averaged over the length of tract) at the first ABCD observation from SEDA intercept while also controlling for age, log-transformed income-to-needs ratio, parental education, family cohesion, neighborhood cohesion, sex, and pubertal status. Family structures nested within scanner site were included as random effects in each model^45^.

Examining the beta-coefficients from these models revealed significant relationships between FA and SEDA intercept in the left and right arcuate, the left posterior arcuate, right cingulate cingulum (CGC), and three colossal tracts. The strongest SEDA relationship between SEDA intercept and FA was observed in the left arcuate (all FDR-corrected p < 0.05; Figure 4, second row from bottom). All together, these tract-wise models demonstrate that the school a child attends influences the development of some white matter tracts above and beyond the myriad of other socioeconomic variables that characterize a child’s environment.

However, as expected, FA in many tracts was also related to other environmental, developmental, and demographic factors (see Figure 2 for overview of these relationships). Parental education was linked to FA in the bilateral arcuate, CST, IFOF, ILF, right VOF, and the right SLF, whereas pubertal status was negatively related to FA in a collection of calossal bundles. Furthermore, age was positively related to higher FA across the entirety of the white matter and males, on average, demonstrated lower FA compared to females across most white matter tracts, with the exception of the left and right CGC and temporal portion of the corpus callosum. These results illustrate that, although educational environment is uniquely linked to FA in some tracts, other environmental and developmental factors are also related to white matter tissue properties across the white matter.

**Figure 2:**
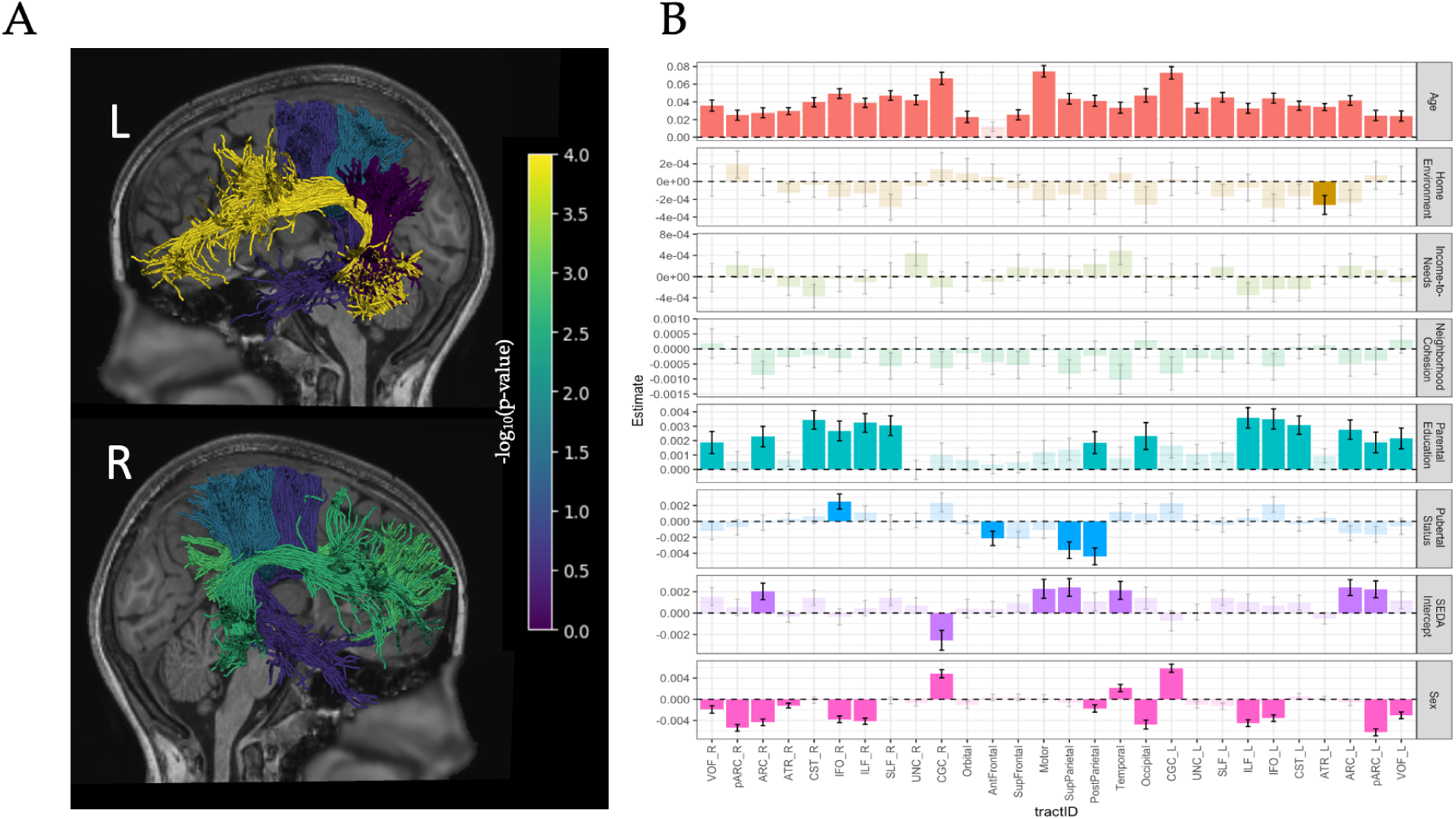
**A.** Renderings of the six white matter tracts significantly related to SEDA intercept. These include the left and right arcuate fasciculus, the left posterior arcuate, and the motor, superior parietal, and temporal bundles of the corpus callosum. Shading represents the -log_10_(p-value) for the beta-weight on SEDA intercept from the models predicting FA in each tract (1.301 corresponds to a p-value of 0.05). This association was strongest in the left arcuate (yellow in the top panel). **B.** Beta-coefficients for the fixed effects of the models predicting FA in each major white matter tract. Each row in the figure refers to the fixed-effect in each model and each color represents a specific bundle. Error bars represent the standard error of each beta-coefficient. Bars that in bold illustrate the beta-weights with FDR-corrected p<0.05.

#### Development of the left and right arcuate is moderated by educational opportunity

Based on the observed cross-sectional link between FA and SEDA intercept in both the left and right arcuate, we then focused on the longitudinal development of these two white matter tracts. Because past longitudinal studies have linked the development of left arcuate and gains in reading skill^28,44,53^, we chose to center this analysis around the left arcuate and its right hemisphere counterpart. To investigate the developmental dynamics of these two white matter tracts, we constructed a series of linear growth models^54^ predicting mean FA over time (operationalized as years since initial MRI scan). In our growth models, we again included individuals nested within family structures nested within scanner sites as random effects. To control for known developmental, demographic, and socioeconomic effects, we also included initial age, pubertal status, sex, log-transformed income-to-needs ratio, parental education, family cohesion, and neighborhood cohesion, sex as fixed-effects. Our main predictors of interest in these models were time, SEDA intercept, tract (right or left arcuate), and their interactions.

We began by fitting separate growth models predicting mean FA in the left and right arcuate. These models revealed significant changes in FA in both tracts within each participant across the two observations (both p < 0.001; See Supplemental Materials for full model outputs). These models also revealed a significant relationship between SEDA intercept and mean FA across both tracts, suggesting that, on average, individuals with greater educational opportunities have higher FA in both the left and right arcuate. We then tested the hypothesis that educational opportunity relates to interindividual differences in FA development over time. To do so, we added a SEDA intercept by time interaction to both growth models. Wald tests comparing the full and reduced models revealed that the addition of the interaction term significantly improved the fit for model predicting FA in the left arcuate (χ^2^(1) = 35.603, p < 0.001) but not the right arcuate (χ^2^(1) = 1.773, p = 0.183), suggesting that educational opportunity may have developmental effects specific to the left arcuate.

To directly test the hypothesis that educational opportunity is related to differences in FA development between the left and right arcuate, we then constructed a combined growth model based on data from both the left and right arcuate fasciculus. This model included the same fixed and random effects as the previous growth models with the addition of a three-way hemisphere by SEDA intercept by time interaction, allowing us directly compare FA development across the left and right arcuate. This model revealed that, on average, FA was lower in the right arcuate compared to the left (B = -0.023; p < 0.001) and that FA increased over time in both tracts (B = 0.002; p < 0.001). Furthermore, the average rate of FA development was slightly, yet significantly, higher in the right arcuate compared to the left (B = 0.0004; p<0.001). Interestingly, individuals in environments with higher SEDA intercept scores, on average, demonstrated significantly faster rates of FA development across both the right and left arcuate (B = 0.001; p < 0.001) and that this interaction was significantly more pronounced in the left arcuate compared to the right (B = -0.0013; p < 0.001; Figure 3). For a full summary of the longitudinal growth model, see Supplementary Table 1.

**Figure 3:**
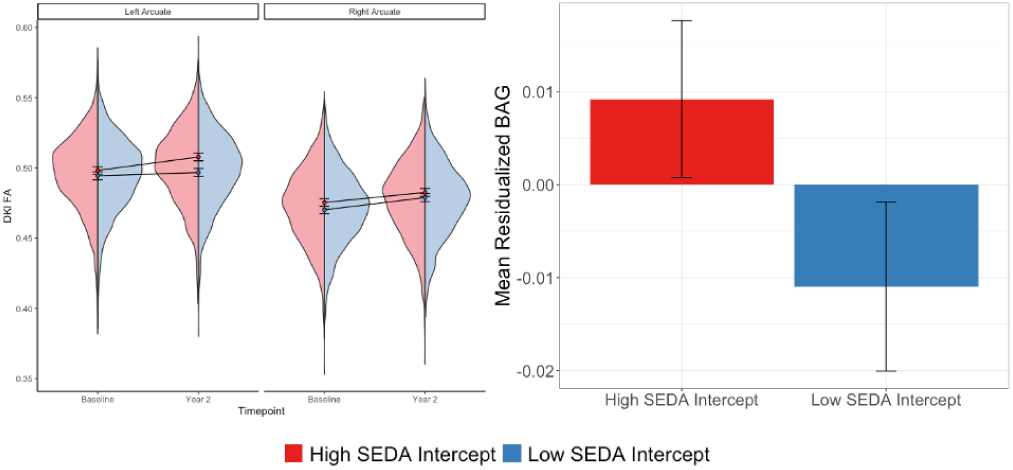
**Left:** Growth trajectories for Diffusion Kurtosis (DKI) FA in the left and right arcuate across the first two observations of the ABCD study. The red and blue lines represent the average DKI FA growth trajectories for individuals in high (Intercept = 1) or low SEDA (Intercept = -1) intercept schools, respectively. Gray lines represent the observed changes in FA in the left and right arcuate for each individual present in the dataset. **Right:** Mean residual values for the model predicting Brain-Age Gap from a reduced model that excludes SEDA intercept as a predictor, but retains all other random and fixed-effects. Each bar represents either the top (red) or bottom (blue) 20% of participants based on their SEDA intercept scores. Error bars represent one standard error from the mean.

### Educational Opportunity Accelerates White Matter Development

Although the bundle-wise analyses suggest that educational opportunity is related to white matter development in the left and right arcuate, cingulate cingulum, and corpus callosum, linear-mixed effects models cannot capture these relationships while considering the entirety of the white matter. To better understand the link between educational opportunity and global white matter development, we trained a series of convolutional neural networks to predict an individual’s age from the white matter properties derived using pyAFQ (see Methods for overview of training procedure). Our brain-age model trained on white matter data from both timepoints, was able to explain roughly 22% of the variance in age in the validation set with an MAE of 0.834 years. It should be noted that the variance explained by this model is smaller than other brain age models ^55^ due to the restricted age range in the ABCD sample. Nevertheless, the residuals from this model, or the difference between the model’s predicted age and each participant’s observed age, can be thought of as the brain-age gap (BAG), a relative measure of how accelerated or delayed an individual’s brain is maturing.

After training our brain age model, we fit a sequence of linear-mixed effects models predicting interindividual differences in BAG from a range of environmental and demographic factors. Our baseline model included age, log-transformed income-to-needs ratio, parental education, family cohesion, neighborhood cohesion, sex, and pubertal status as fixed-effects (and the same random effects structure as previous models), allowing us to control for known demographic, socioeconomic, and developmental factors related to brain development^19,56^.

To test the relationship of educational opportunity and white matter development, we fit a model including SEDA intercept as an additional predictor, and compared the full and reduced models using a Wald test. This tests revealed that SEDA intercept significantly improved model fit (χ^2^(1) = 5.483, p = 0.02). The coefficients of the SEDA intercept model revealed a significant negative relationship between BAG and age (B=-0.475; p<0.001; See Supplemental Table 2 for full model output), suggesting that our model underestimates the brain-age of older participants and over estimates the brain age of younger participants. This is a known phenomenon with brain age models^57^ and can be interpreted as regression to the mean. Additionally, the model revealed significant relationships between BAG and both parental education (B=0.038; p=0.02) and pubertal status (B=0.0041; p<0.001). The effect of parental education is in line with past findings^13,16,18^ suggesting that parental education is linked with differences in white matter development and the effect of pubertal status suggests that puberty accelerates white matter development.

Interestingly, this model also revealed a small, yet significant relationship between the BAG and SEDA intercept (B=0.019; p=0.019; Figure 3). The observed relationship between SEDA intercept and BAG suggests that the educational opportunities afforded to a learner before 3^rd^ grade are linked to global differences in white matter maturation. These results, coupled with the insignificant coefficients on other demographic and environmental predictors (all p>0.05), indicate that an individual’s early educational opportunities are related to global white matter development even when accounting for factors such as income-to-need ratio, family cohesion, and neighborhood stability.

## Discussion

In the present study, we leveraged the unique epidemiological sample from the ABCD study to explore the relationship between an individual’s white matter development and the educational opportunities provided by their early childhood and elementary school environments. The scale of the ABCD study, coupled with the rich demographic measures present in the data, provides the very first opportunity to examine how the diverse educational experiences found across the United States’ educational system relate to brain development, while also accounting for other environmental factors. We leveraged data from the Stanford Education Data Archive alongside diffusion MRI data to test the hypothesis that an individual’s white matter development is related to the quality of their educational environment.

These analyses showed that SEDA intercept, a measure of early educational environment, was associated with fractional anisotropy in the left and right arcuate fasciculi, left posterior arcuate, and the corpus callosum. These relationships held even when accounting for other socioeconomic factors known to relate to white matter, such as parental education or household income. When examining the longitudinal relationship between white matter properties and educational opportunity over the course of 2 years, the rate of FA development in both the right and left arcuate was associated with SEDA intercept and this relationship was slightly, yet significantly, more pronounced in the left arcuate. Furthermore, a global analysis of the white matter using a brain-age modeling approach revealed a relationship between SEDA intercept and levels of white matter maturation.

To the best of our knowledge, these results are the first to show a specific link between educational environment and white matter development in a sample of this size. As observed in the present study, measures of socioeconomic status are highly correlated with one another. Thus, some indices of SES, such as household income, may confound other measures of SES in studies of brain development. The fact that SEDA intercept predicts tissue properties of specific white matter tracts, as well as global brain-age measures, even when controlling for other developmental and environmental factors, demonstrates a specific relationship between an individual’s educational environment and their white matter development.

Because SEDA intercept can be thought of as a measure of the educational opportunities available to a learner in early childhood and elementary school^43,58^, these results suggest that early educational experiences impact the development of white matter tracts throughout elementary school and into middle school. This parallels behavioral and educational policy research that has shown that gaps in reading and mathematics at the onset of elementary school, on average, persist throughout the course of K-12 education^59–61^ and that early measures of academic skills serve as strong predictors of later academic success and life outcomes^5,6^.

However, it remains unclear whether early childhood opportunities continue to influence white matter development throughout late elementary school and into adolescence or if the year-to-year educational opportunities afforded by a school also shape white matter later in development. Future work with the full longitudinal ABCD sample will have to explore this question. Nevertheless, the findings that SEDA intercept is related to both increased rates of FA development in the left arcuate and overall brain-age indicate that differences in early educational opportunity may not only lead to differences in academic outcomes but also influence the developmental trajectories of the white matter throughout childhood and into adolescence.

Furthermore, the observed relationship between SEDA intercept and the brain-age gap suggest that early educational opportunities may not only influence the development of the specific white matter tracts underlying academic skills, but rather relate to white matter development more broadly throughout the brain. Studies in animal models have shown that environmental enrichment leads to an increase in cellular activities related to myelination, such as the proliferation of oligodendrocyte progenitor cells and alterations of the oligodendrocyte translatome in a broad range of brain regions^62–64^. Furthermore, evidence from human neuroimaging data suggests that environmental stress and caregiving settings relate to differences in white matter properties throughout the brain^65,66^. Taken together, these results suggest that general enrichment of an individual’s educational environment may drive global changes in white matter, whereas opportunities to meaningfully engage in specific subject areas may impact the white matter tracts subserving academic skills.

Unfortunately, because SEDA scores are derived from school-level standardized test scores, they do not provide any insight into specific aspects of each individual’s learning experience within a given environment, thereby limiting the specificity of the questions we can answer using the ABCD dataset. A student’s experience in the classroom can be impacted by socio-cultural equity, language use, student-teacher relationships, the curriculum adopted by the school district, and classroom organization^67–70^, all of which do not necessarily manifest in their school’s standardized test scores. These factors may differentially impact global white matter development and tissue properties in white matter connections subserving different academic skills. Future intervention studies and additional measures of educational environment are needed to better understand these relationships. These will serve to isolate the white matter changes driven by opportunities to engage in an academic subject area, such as reading or mathematics, from those due to aspects of the educational environment that are independent of the specific academic content matter.

Furthermore, because each participant has at most two observations, our longitudinal models are limited in the types of relationships captured by difference scores. White matter properties have been shown to follow non-linear growth trajectories over the lifespan^66,71^, however, with only two observations, one cannot effectively model non-linear relationships. Future research using the full longitudinal ABCD sample will have to explore the developmental dynamics of the white matter over the course of adolescence and determine whether the observed relationship between FA development and SEDA is best described by a linear or non-linear trajectory.

In summary, these results suggest that the educational opportunities provided to a learner in early elementary school are related to subtle differences in white matter maturation, even when accounting for other socioeconomic factors. We observe a brain-wide link between white matter development and educational environment and find that this relationship is strongest in the white matter tracts typically associated with academic skills. Future research is needed to inform the design of interventions and policies addressing educational inequities from a neuroscientifically-informed perspective. The current study provides the first direct evidence for the relationship between educational opportunity and brain development at scale and shed light on the complex interaction between environmental factors, brain development, and learning.

## Methods

### Participants

The participants in the present study come from the ABCD study, a ten-year longitudinal study that includes both neuroimaging and behavioral data collected from children aged 9-10 from 21 study sites across the United States^38^. The data used in the present analysis come from the baseline and 2-year follow up visits of the ABCD study and can be found in the ABCD curated annual data release 4.0 (https://nda.nih.gov/abcd/). The baseline observation included 6,410 individuals who had the necessary neuroimaging and demographic data and the longitudinal data included 4,770 individuals with the necessary data at both time points.

### Covariates of Interest

We included a range of demographic and developmental factors as covariates including participant age, log-tranformed income-to needs ratio, parental education, family cohesion, neighborhood cohesion, biological sex, and pubertal status. All of these measures are readily available or calculated using the data present in the ABCD data release 4.0.

### Income-to-Needs Ratio

Log-transformed income-to-needs ratio was calculated using the approach outlined in Wiessman et al. (2023), which combines family income and household size data. Briefly, in the ABCD study, parents report family income on a scale of 1-10, where each interval represents an income range. The midpoint of the reported range was then calculated for each participant. This dollar amount was then divided by the poverty threshold for a household of a given size. The thresholds used in this calculation come from the 2017 report by the U.S Census Bureau^72^. This value was then log-transformed.

### Parental Education

The ABCD study records parental reports of the highest level of education they have completed. This is measured on an ordinal scale ranging from “Never attended school” to “Doctoral Degree”. In the present analysis, we converted this ordinal scale into a binary variable indicating whether or not a participant’s parent completed any sort of post-secondary education (associates degree/bachelors degree).

### Non-Academic Environment

Family home environment was measured using the average of the nine questions present on Family Environment Scale-Family Conflict^73^. A higher score on this measure indicates higher levels of conflict within an individual’s household and family environment. Neighborhood cohesion was assessed by taking the average of the ten items present on the ABCD Parent PhenX Community Cohesion measure^74^. For this measure, a higher score indicates that the participant perceives their neighborhood and surrounding community as safer and more cohesive.

### Pubertal Status

Pubertal status was assessed using the PDS^75^, a measure designed to mimic the Tanner scale to assess the development of secondary sex characteristics during the onset of puberty. In line with past research using PDS in the ABCD sample^56,76^, pubertal status was calculated by taking the average of the seven PDS items present on the parental PDS survey collected at each time point.

### Educational Opportunity

Educational opportunity was measured using linked data from the Stanford Education Data Archive (SEDA)^27^. This dataset leverages standardized test scores from 3rd to 8th grade students in nearly every single school district in the United States to generate measures of the educational opportunities provided by a given school or district relative to the national average. The SEDA database breaks educational opportunity into two distinct measures: intercept and slope. SEDA intercept refers to the average standardized test score for third graders from a given school or district relative to the national average and can be thought of as an index of the pre-school and early elementary school educational opportunities provided by a school catchment area^43^. SEDA slope is a measure of year-to-year growth in standardized scores in students from a given school or district relative to the national average. This can be thought of as the educational opportunity provided to students in a school or district between 3rd and 8th grade.

### Diffusion MRI Acquisition and Processing

The neuroimaging data used in this analysis come from the baseline and Year 2 follow-up sessions collected across the 21 ABCD study sites. An overview of the data acquisition and preprocessing protocols can be found in Casey et al. (2018)^38^ and Hagler Jr. et al. (2019)^77^. Briefly, multi-shell, high angular-resolution imaging scans were collected on each participant during each scan session. These data underwent manual quality control and were then minimally preprocessed using a pipeline that included eddy-current correction, motion correction, *B*_0_ distortion correction, and gradient warp correction^77^.

These preprocessed diffusion images were then processed with pyAFQ^41^. Briefly, fiber orientation distributions were estimated in each voxel using constrained spherical deconvolution^78^ implemented in DIPY^79^ before probabilistic tractography was used to generate streamlines throughout the white matter. As originally described in Yeatman et al. (2012)^42^, 30 major white matter tracts were identified from these streamlines. Each tract was then sampled to 100 nodes. At each node, fractional anisotropy (FA), mean diffusivity (MD), radial diffusivity (RD), and axial diffusivity (AD) were calculated using the diffusion kurtosis model (DKI)^80,81^.

To account for potential non-biological variance in the diffusion MRI signal introduced by scanner differences across the 21 ABCD sites, ComBat harmonization^46,47,82^ was performed on the diffusion metrics calculated by pyAFQ. Harmonization was performed using the neurocombat_sklearn Python library^47,82^.

### Brain Age Gap Analysis

To generate age predictions for the brain-age gap analysis, we trained a ResNet model^83^ implemented in the Python library AFQ-Insight^55,84^ on the harmonized pyAFQ outputs from both the baseline and Year 2 follow-up scans. The data used to train and evaluate this model were split into three splits: a training set, a test set, and a validation set. To prevent peeking, longitudinal observations from the same participant were placed in the same split. The validation set contained 20% of the observations, while the remaining 80% was distributed across the training and test sets. To prevent overfitting, the model was then trained on varying proportions of the training and test sets, allowing us to determine the point at which the model performance did not improve with the addition of more training data (Supplemental Figure 2).

We found that model performance plateaued when the model was trained on 56% of the overall sample. This model attained an R^2^ score of 0.22 on the unseen validation set. To prevent data leakage in our brain age models, we then generated two additional train-test splits to ensure that brain age predictions for each individual were generated from a model that was trained on data that did not include that individual. We then trained two additional models on 56% of the overall sample to generate brain age predictions for the individuals used as the training set of the initial brain age model. Across these three models, the average R^2^ score was 0.19 on the unseen data. We then calculated the residuals of these brain age predictions for each individual, using the prediction from the model that was not trained on that individual’s data. The residuals of these brain age model were then used as the outcome measure of a linear-mixed effects model to explore the relationship between educational opportunity and brain-age, while controlling for other developmental and socioeconomic factors. All linear-mixed effects models were carried out with R version 4.2.1^85^ using the *lme4* package (version 1.1.30)^86^.

### Growth Modeling

To investigate intraindividual change in the white matter properties of the left and right arcuate and the relationship of this change with educational opportunity, we constructed a series of growth models^54^ specified as follows:

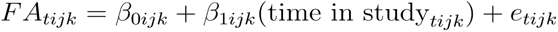

where each participant’s FA at a given scan session, *t*, is modeled as a function of a participant specific intercept (β_0i_), a participant specific slope (β_1i_), and a residual error term (*e_ti_*). To examine interindividual differences, the participant specific coefficients were modeled as:

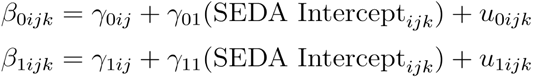

where the γ coefficients on SEDA Intercept refer to, on average, how baseline FA and FA development differ with SEDA intercept and *u_0jki_* and *u_1ijk_* refer to residual error at the individual level. These models also included initial age, log-transformed income-to-needs ratio, parental education, family home environment, neighborhood cohesion, and pubertal status as covariates.

Our final growth model, which examined FA development in the left and right arcuate simultaneously, included two additional parameters, γ_02_(Hemisphere_ijk_) and γ_12_(Hemisphere_ijk_), which allowed us to compare FA development across the two tracts. In this model γ_0*ijk*_ and γ_0ijk_ are modeled at the family structure level as:

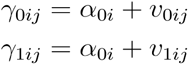

where *v_0ij_* and *v_1ij_* refer to residual error at the family structure level and α_00_ and α_01_ are modeled at the level of scanner site as:

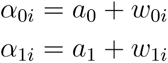

where *w_0i_* and *w_1i_* refer to residual error at each scanner site and α_0*i*_ and α_1*i*_ are the mean FA and rate of FA development, respectively, at each scanner site.

These models were fit in R version 4.2.1^85^ using the *lme4* package (version 1.1.30)^86^. Code to replicate the analyses and figures presented in this manuscript can be found at: https://github.com/earoy/white_matter_education

---

**Supplemental Figure 1:**
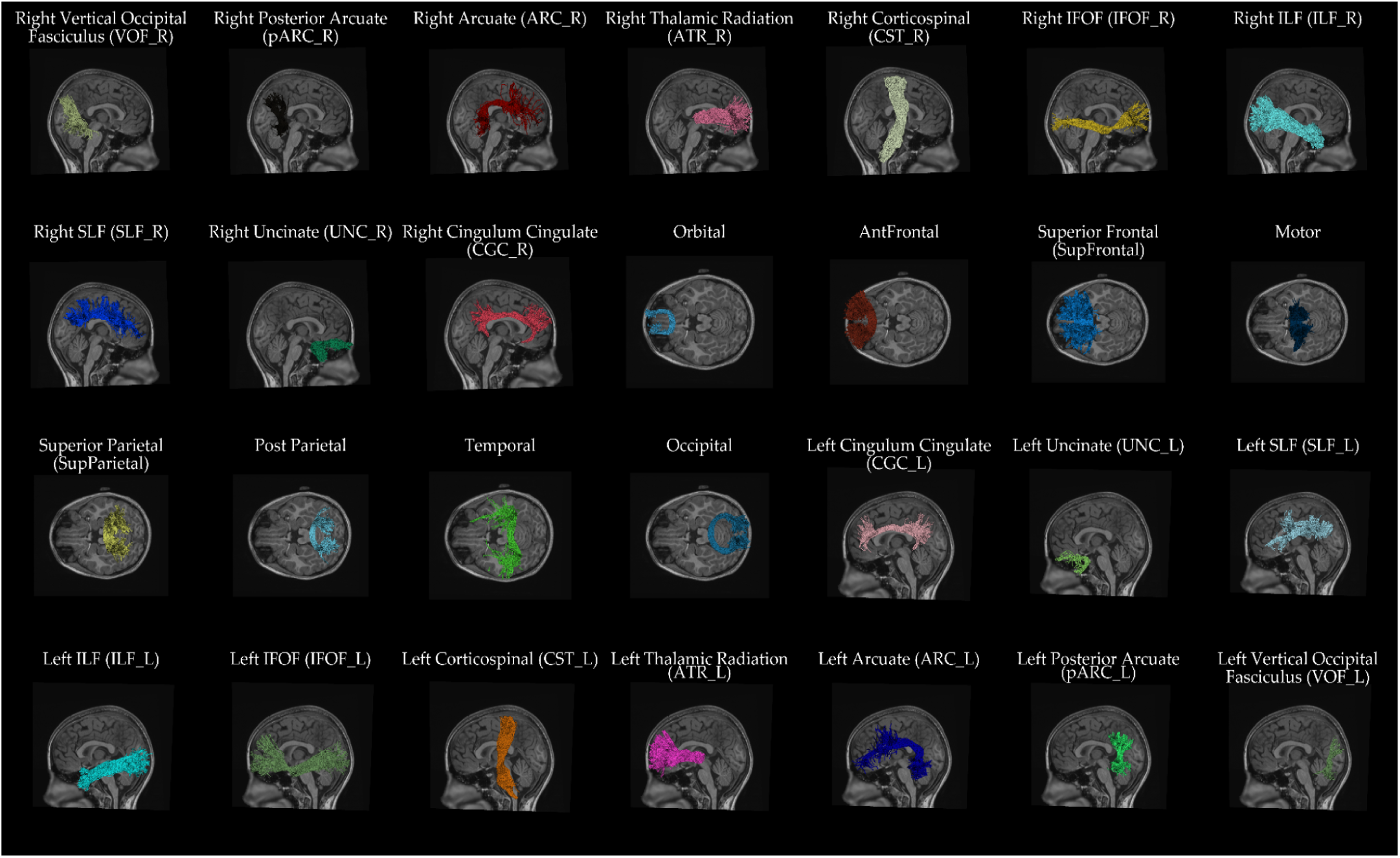
Bundle renderings of the 28 white matter tracts identified by pyAFQ

**Supplemental Figure 2:**
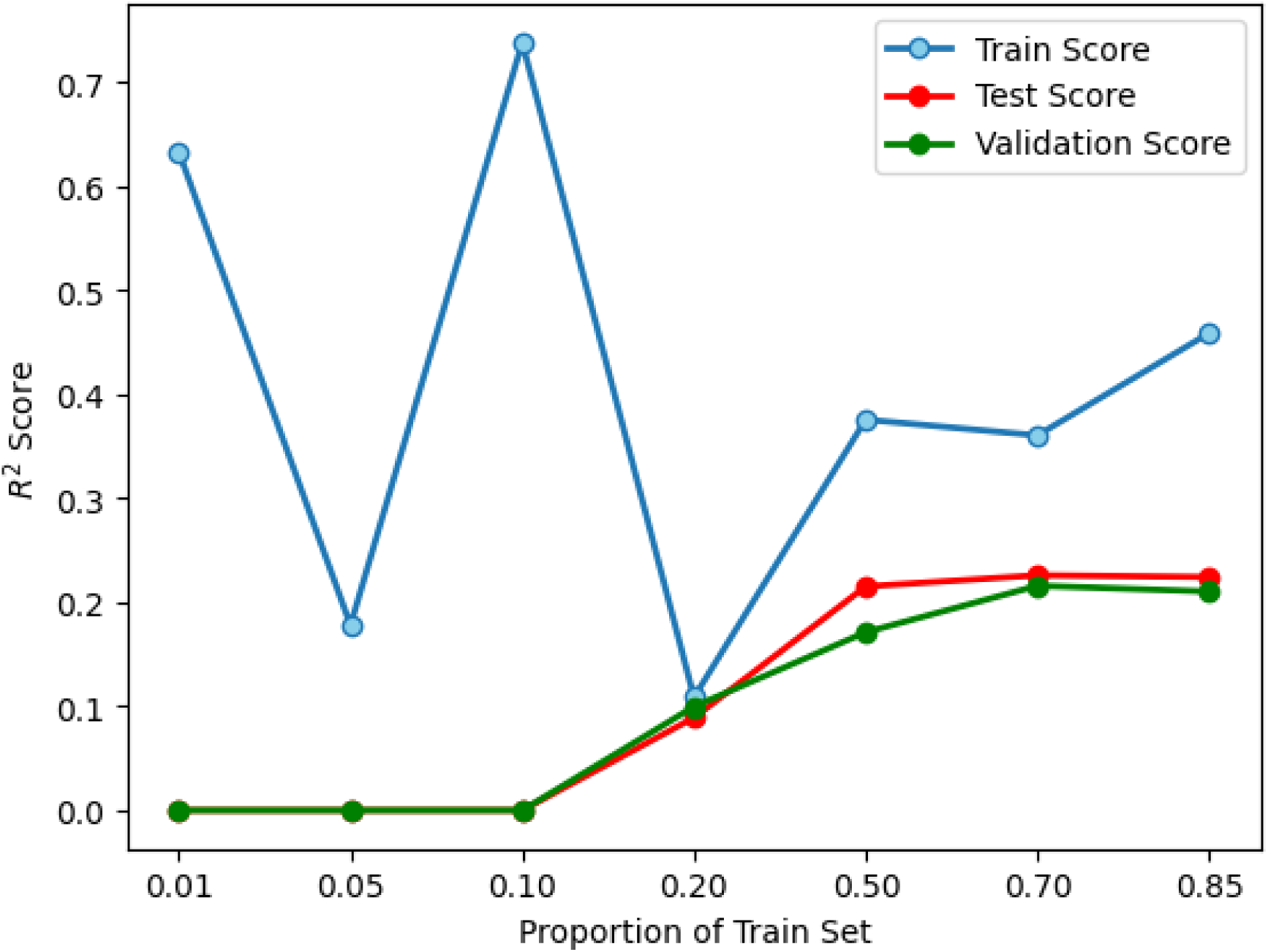
An overview of brain-age model performance trained on varying proportions of the test split. The x-axis represents the proportion of the training set used to train the brain age model and the y-axis represents the R^2^ score of each model.

**Supplemental Table 1:**
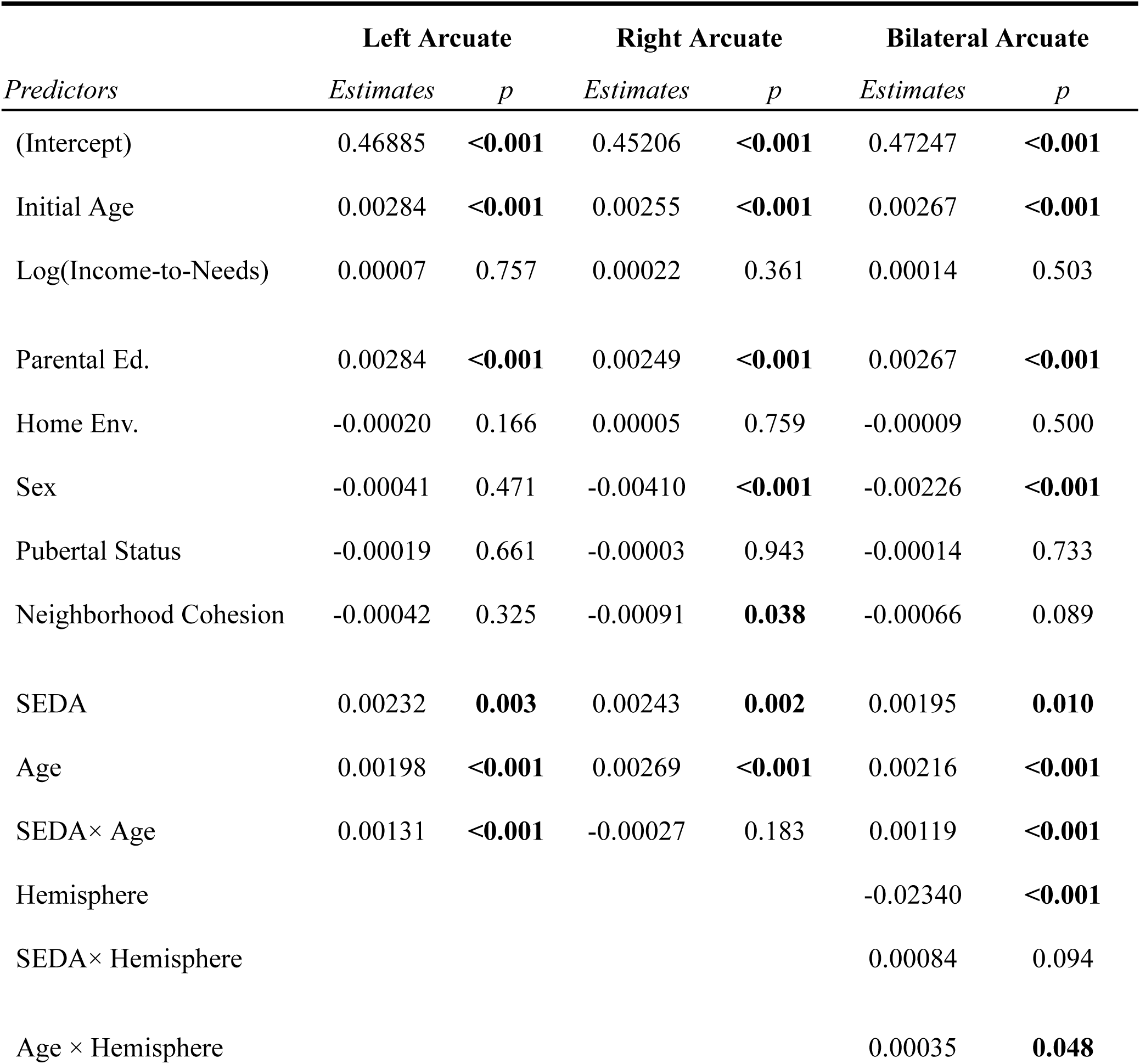
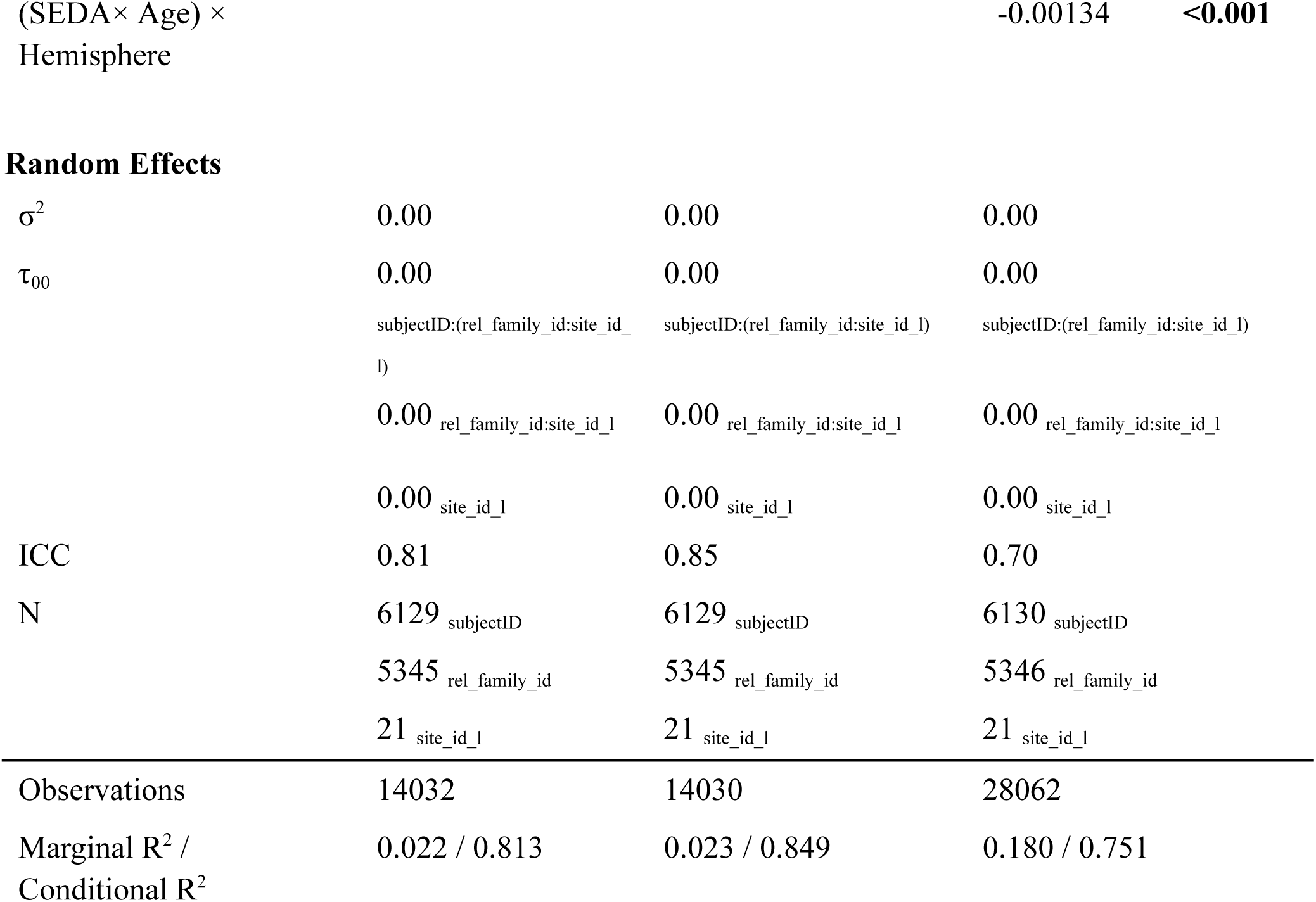
Output for the growth model predicting FA development in the left arcuate, right arcuate, and left and right arcuate together. In the bilateral model, the left arcuate is used as the reference predictor.

**Supplemental Table 2:**
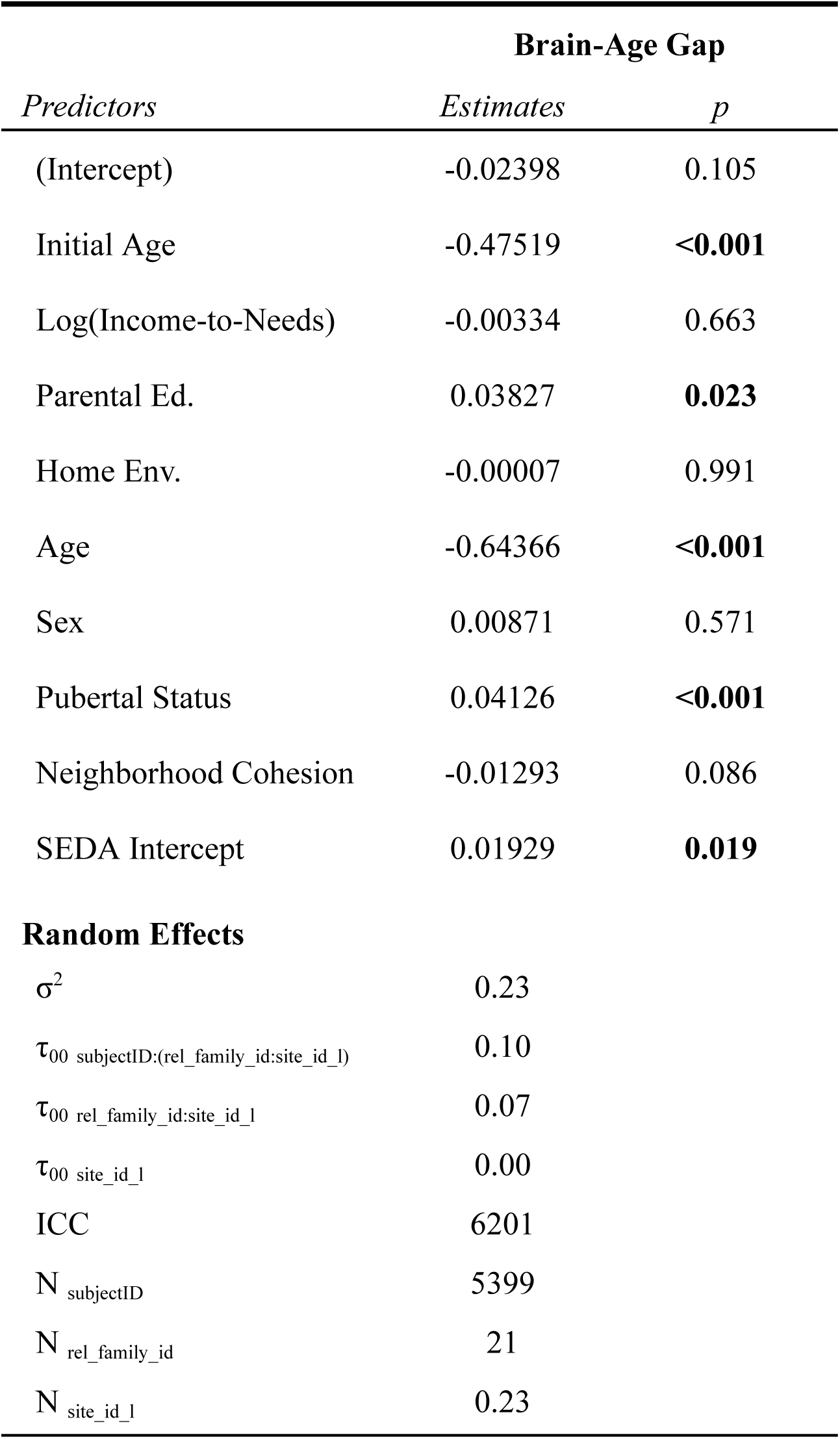
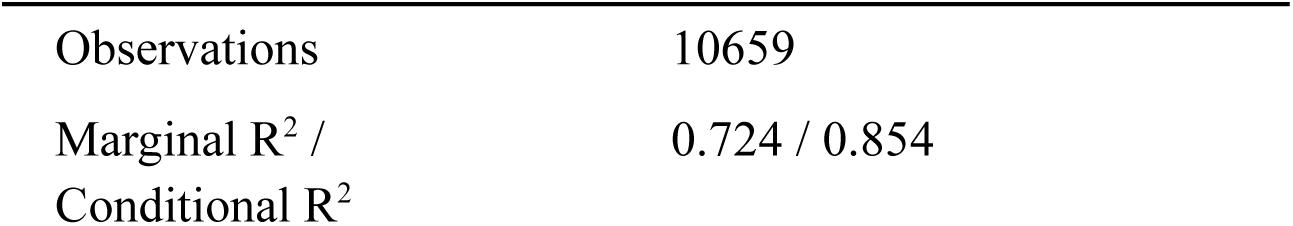
Output for the growth model predicting brain-age gap.

| **Brain-Age Gap** |  |  |
| --- | --- | --- |
| *Predictors* | *Estimates* | *p* |
| (Intercept) | -0.02398 | 0.105 |
| Initial Age | -0.47519 | **<0.001** |
| Log(Income-to-Needs) | -0.00334 | 0.663 |
| Parental Ed. | 0.03827 | **0.023** |
| Home Env. | -0.00007 | 0.991 |
| Age | -0.64366 | **<0.001** |
| Sex | 0.00871 | 0.571 |
| Pubertal Status | 0.04126 | **<0.001** |
| Neighborhood Cohesion | -0.01293 | 0.086 |
| SEDA Intercept | 0.01929 | **0.019** |
| **Random Effects** |  |  |
| $\sigma^2$ | 0.23 | |
| $\tau_{00}$ subjectID:(rel_family_id:site_id_1) | 0.10 | |
| $\tau_{00}$ rel_family_id:site_id_1 | 0.07 | |
| $\tau_{00}$ site_id_1 | 0.00 | |
| ICC | 6201 |  |
| N<sub>subjectID</sub> | 5399 |  |
| N<sub>rel_family_id</sub> | 21 |  |
| N<sub>site_id_1</sub> | 0.23 |  |

|  |  |
| --- | --- |
| Observations | 10659 |
| Marginal R<sup>2</sup> /<br>Conditional R<sup>2</sup> | 0.724 / 0.854 |

